# Individual-level specialisation and interspecific resource partitioning in bees revealed by pollen DNA metabarcoding

**DOI:** 10.1101/2021.08.01.454648

**Authors:** Jan Klečka, Michael Mikát, Pavla Koloušková, Jiří Hadrava, Jakub Straka

## Abstract

It is increasingly recognised that intraspecific variation in traits, such as morphology, behaviour, or diet is both ubiquitous and ecologically important. While many species of predators and herbivores are known to display high levels of between-individual diet variation, there is a lack of studies on pollinators. It is important to fill in this gap because individual-level specialisation of flower-visiting insects is expected to affect their efficiency as pollinators with consequences for plant reproduction. Accordingly, the aim of our study was to quantify the level of individual-level specialisation and foraging preferences, as well as interspecific resource partitioning, across different temporal scales in three co-occurring species of bees of the genus *Ceratina* (Hymenoptera: Apidae: Xylocopinae), *C*. *chalybea*, *C*. *nigrolabiata*, and *C*. *cucurbitina*. We conducted a field experiment where we provided artificial nesting opportunities for the bees and combined a short-term mark-recapture study with the dissection of the bees’ nests to obtain repeated samples from individual foraging females and complete pollen provisions from their nests. Hence, we could study variation of the composition of pollen collected by the bees at different temporal scales. We used DNA metabarcoding based on the ITS2 locus to identify the composition of the pollen samples. We found that the composition of pollen carried on the bodies of female bees and stored in the brood provisions in their nests significantly differed among the three co-occurring species. At the intraspecific level, individual females consistently differed in their level of specialisation and in the composition of pollen carried on their bodies and stored in their nests. Our study thus provides evidence of consistent individual-level specialisation in pollinators across multiple temporal scales. We also demonstrate that higher generalisation at the species level stemmed from larger among-individual variation in diets as observed in other types of consumers, such as predators. Our study thus reveals how specialisation and foraging preferences of bees change from the scale of individual foraging bouts to complete pollen provisions accumulated in their nests over their lifetime. Such multi-scale view of foraging behaviour is necessary to improve our understanding of the functioning of plant-flower visitor communities.

## INTRODUCTION

Intraspecific variation in morphological and physiological traits, behaviour, and diet is common in many types of animals and has important implications for ecological processes at the population and community levels (Bolnick et al., 2003, 2011; Araújo et al., 2011). For example, between-individual variation in specialisation and dietary preferences may have strong effects on the structure and stability of ecological networks Bolnick et al. (2011). These effects stem from a potential disconnect between specialisation at the species (or population) level and the individual level. Total niche width of a species can be decomposed into individual-level niche width and between-individual variation (Roughgarden, 1972, 1974). An animal may thus be generalised at the species level in two fundamentally different ways: either all individuals have a similar generalised diet or different individuals are specialised on different resources. Although the existence of between-individual variation in diet has been recognised for a long time and formed a basis of Van Valen’s niche expansion hypothesis (Van Valen, 1965), the potential ecological importance of between-individual diet variation has been neglected until a relatively recent resurgence of interest in individual variation (Bolnick et al., 2003; Araújo et al., 2011; Bolnick et al., 2011).

A large amount of evidence demonstrating that many species of animals have high levels of between individual variation in diets has been accumulated (Bolnick et al., 2003; Araújo et al., 2011), but most published studies focused on predators, particularly vertebrates. We are aware of only two studies on flower-visiting insects which studied between-individual variation in diets using repeated observations of the same individuals as required to properly describe individual diets (Bolnick et al., 2002, 2003). In their landmark study, Heinrich (1976) found that individual bumblebees specialised on different flowering plants not only during a single foraging bout, but also over a longer time frame, although the evidence was rather anecdotal. More recently, Szigeti et al. (2019) provided quantitative evidence for between-individual variation in flower visitation by a butterfly, *Parnassius mnemosyne*, partly related to temporal changes in flower abundance. However, more data are needed to test the generality of these results and to evaluate their implications for plant-pollinator interactions (Brosi, 2016).

To make matters more complicated, specialisation may further vary at different temporal scales within an individual, e.g. a pollinator may be highly specialised during a single foraging bout, which is often called “floral constancy” or “flower constancy”, but have a substantially broader diet over its lifetime (Heinrich, 1976; Brosi, 2016). Flower constancy may have a strong effect on the reproductive success of insect-pollinated plants because high specialisation during a single foraging bout increases both male and female fitness of the plants by increasing the probability of pollen transfer to a flower of the same plant species and by minimising the deposition of heterospecific pollen harmful to plant female fitness (Waser, 1978; Morales and Traveset, 2008). Although flower constancy has been demonstrated in many pollinators, including social and solitary bees, butterflies, and hoverflies (Heinrich, 1976; Waser, 1986; Lewis, 1986; Goulson and Wright, 1998; Slaa et al., 1998; Amaya-Márquez, 2009), it considers foraging decisions only over a very short temporal scale, often over only several consecutive flower visits. On the other hand, we lack information on the variation among foraging bouts of the same individuals over a longer time scale with the few exceptions mentioned above (Heinrich, 1976; Szigeti et al., 2019).

Embracing a multi-level view of foraging specialisation, with the partitioning of individual-level specialisation, between-individual diet variation, and overall population- or species-level specialisation, can shed new light on interspecific interactions (Brosi, 2016). So far, it is known that large between-individual variation decreases the strength of intraspecific competition because each individual competes only with a subset of conspecifics, but it may increase the strength of interspecific competition. Species and individuals may thus respond in different ways to changes in the strength of intraspecific or interspecific competition, such as by changing individual diet width (Fontaine et al., 2008; Brosi and Briggs, 2013) or changing the level of diet overlap among individuals in the population (Van Valen, 1965; Bolnick et al., 2007). Different strategies may be employed also by different individuals in the same population. For example, some individuals may be specialised, which makes them more efficient in resource use (Strickler, 1979; Hofstede and Sommeijer, 2006), while others are more generalised. Switching between resources incurs costs because of memory and learning constraints (Lewis, 1986; Gegear and Laverty, 1998), but more flexible individuals capable of switching between different types of resources may better cope with spatial and temporal variation in resource availability (Hofstede and Sommeijer, 2006). At the species level, there is a strong support for the hypothesis that populations with larger between-individual variation are less vulnerable to environmental changes in various groups of organisms (Forsman and Wennersten, 2016). High between-individual variation together with the foraging flexibility of flower-visiting insects may underpin the robustness of plant-flower visitor networks to habitat destruction (Noreika et al., 2019) or loss of resources (Biella et al., 2019a, 2020). Studies combining measures of between-individual diet variation and interspecific resource partitioning are thus needed to shed more light on the ecological consequences of individual-level diet variation.

We studied foraging preferences and specialisation in three sympatric species of mostly solitary bees of the genus *Ceratina* to address several of the current knowledge gaps. Specifically, we used pollen DNA metabarcoding to analyse the level of specialisation in foraging for pollen at the interspecific and intraspecific levels. We compared pollen composition among nests build by different females, individual brood cells within the nests, and pollen collected during individual foraging bouts. Our aim was to test whether the three sympatric species differed in their foraging preferences, diet breadth at the species and individual levels, and in between-individual variation in diet composition. Such differences in foraging strategies could decrease the intensity of resource competition and facilitate species coexistence.

## MATERIALS & METHODS

### Species studied

The genus *Ceratina* Latreille, 1802 (Hymenoptera: Apidae: Xylocopinae) is a cosmopolitan genus of bees whose common ancestor was probably facultatively eusocial (Rehan et al., 2012). Most extant *Ceratina* species are also facultatively eusocial (Groom and Rehan, 2018), but the proportion of social nests is generally low, and solitary nesting prevails particularly in temperate climates (Groom and Rehan, 2018; Mikát et al., 2020a). Also, some species are known for complex parental care (Mikát et al., 2016), including the only known example of biparental care in bees (Mikát et al., 2019). We focused on three species of the genus *Ceratina*, which are the most abundant bee species at the study site (see below), namely *C*. *chalybea*, *C*. *nigrolabiata*, and *C*. *cucurbitina*. All three species are morphologically similar and live mostly in warm grassland habitats. They build their nests in dead plant stems with soft pith, e.g. of *Rosa canina*, *Centaurea* spp., *Verbascum* spp., etc. This makes it easy to study their nesting behaviour and obtain pollen samples from their nests. The nest is made of a linear sequence of brood cells, whose relative age is easy to determine: the innermost brood cell is the eldest (Rehan and Richards, 2010). Although biparental care has been observed in *C*. *nigrolabiata*, only the female provisions the nest, while the male’s role is to guard the nest (Mikát et al., 2019). Hence, only a single female provisions each nest during the brood establishment in the species we studied.

### Study site and experimental design

We conducted a field experiment in the Havranické vřesoviště Natural Monument, in the Podyjí National Park, near Znojmo, in the Czech Republic (GPS: 48.8133N, 15.999E) in the spring and summer 2017. The administration of the Podyjí National Park provided a research permit. The study site and its surroundings comprises of a heathland and dry open grasslands with solitary trees and shrubs. We installed artificial nesting opportunities in the grassland following the methods used in previous research at the study site (Mikát et al., 2016, 2019). The artificial nesting opportunities consisted of sheaves containing 20 cut stems of *Solidago canadensis*. Each stem was 40 cm long. The sheaves were attached in a vertical orientation to a thin bamboo stick fixed to the ground. We distributed several hundred sheaves as nesting opportunities in the study site in April before the beginning of the nesting season.

### Sampling in the field

Field sampling consisted of two phases. In the first phase, we collected a subset of the occupied artificial nests on 5 July 2017 and sampled all pollen stored in individual nests. We collected the nests after the end of the bees’ foraging activity, around sunset, when the female bees can be usually found inside the nests (Mikát et al., 2016, 2017, 2019), which allowed us to reliably identify the species to which the nest belonged. We carefully opened the nests in a field laboratory with clippers and collected the pollen provisions from individual brood cells using sterilised forceps, stored the samples in individual microtubes, and dried them at room temperature in a desiccator with silica gel. The ID of the nest and the ID of the brood cell within the nest (brood cells ordered as 1, 2, etc. starting with the eldest one) was recorded along with information about the developmental stage of offspring in each brood cell (egg, larva with its instar identified, or pupa), which we use to estimate the relative brood cell age. Most of the nests were not yet fully developed, i.e. they contained mostly eggs and larvae, only some of them contained pupae in the oldest brood cells, and no offspring has matured yet. We collected pollen from brood cells with unconsumed provisions, i.e. those containing eggs or young larvae (alive or dead). In total, we obtained 227 samples from 66 nests of these three species containing a sufficient amount of pollen for the purpose of our analyses (i.e., unconsumed pollen in at least two brood cells); 52 samples from 17 nests of *C*. *chalybea*, 131 samples from 36 nests of *C*. *nigrolabiata*, and 44 samples from 13 nests of *C*. *cucurbitina*.

In the second phase of the fieldwork, we conducted a mark-recapture study of the three *Ceratina* species from 29 July to 1 August 2017. We used the same type of artificial nests as described above arranged in an array over the area of ca. 10 × 5 m. We individually marked females of the three species captured during the provisioning of their nests. The females were marked by a combination of colour spots on the abdomen. Females were recaptured during four days when they were returning to their nests from foraging bouts. This allowed us to sample pollen collected by the captured female during individual foraging bouts. Capturing the females on return to their nests was facilitated by blocking the entrance to their nests while they were foraging (Mikát et al., 2017). We used sterile aspirators for individual recaptures to prevent contamination. We briefly anaesthetised the captured bee using CO_2_, scrapped the pollen carried on the underside of the abdomen using a single-use toothpick with a small piece of cotton attached to the end (a miniature analogue of cotton buds for ear cleaning), and stored the pollen in a 2 ml tube. In total, we collected 67 samples; 26 samples from 17 females of *C*. *chalybea*, 35 samples from 23 females of *C*. *cucurbitina*, but only six samples from five females of *C*. *nigrolabiata*.

### Pollen DNA metabarcoding

We extracted DNA from the pollen samples using the Macherey-Nagel NucleoSpin Food kit (Macherey-Nagel, Dűren, Germany) according to “the isolation of genomic DNA from honey or pollen” supplementary protocol developed by the manufacturer. Prior to DNA extraction, we homogenised each pollen sample with the CF Buffer from the NucleoSpin Food kit in a 2 ml tube using ceramic beads in a Precellys homogeniser similarly to Bell et al. (2017).

We amplified the ITS2 region (Chen et al., 2010) using standard primers for plant ITS2 used also in previous studies on pollen metabarcoding (Sickel et al., 2015; Bell et al., 2017). Our DNA metabarcoding strategy followed general recommendations by Taberlet et al. (2018). We performed three independent PCR replicates for each sample. The primer design incorporated 8 bp long tags in both the forward and reverse primer, which allowed us to tag individual PCR replicates of individual samples by a unique combination of tags on the forward and reverse primers. The PCR replicates were thus tagged, sequenced together in a single sequencing library, and analysed separately. We used three types of controls: blanks, PCR negative controls, and PCR positive controls. We used a mixture of DNA extracts of five exotic plant species as the PCR positive control. We did the PCR in strips rather than plates to limit cross contamination (Kitson et al., 2019). Each strip contained seven samples and one of the controls. In total, we had 39 blanks, 39 PCR negative controls, and 36 PCR positive controls. The extensive use of different types of controls allowed us to evaluate different sources of contamination and sequencing errors during data analysis (De Barba et al., 2014; Taberlet et al., 2018). PCR cycles included an initial period of 3 min at 95°C; followed by 35 cycles of 30 s at 95°C, 30 s at 55°C, and 1 min at 72°C; followed by a final extension of 10 min at 72°C as in Bell et al. (2017), which seemed to be ideal parameters based also on our preliminary tests. We verified the success of PCR using gel electrophoresis prior to library preparation.

We pooled equal volume of the PCR product from all samples and purified the resulting amplicon pool using magnetic beads (Agencourt AMPure PCR purification kit). The final amplicon pool had a concentration of 52 ng/μl measured by Invitrogen Qubit 3.0 fluorometer (Thermo Fisher Scientific). Library preparation was done using a PCR-free approach with Illumina adaptors added by ligation at Fasteris (Switzerland) and the library was sequenced on Illumina HiSeq 2500 Rapid Run, using 1/10 of the capacity of one sequencing lane resulting in 35,121,401 raw paired reads.

### Reference plant database

We assembled a detailed reference database of ITS2 sequences of most plant species growing in the vicinity of the study site. We attempted to obtain an exhaustive list of plant species growing within the radius of at least 1 km around the study site by our own botanical survey and by extracting data from the literature, particularly a detailed atlas of plants of the Podyjí National Park (Grulich, 1997) and a national database of plant records (Wild et al., 2019). We collected tissue samples (usually leaves) of most entomogamous plant species we could find in the field and identify reliably and dried them with silica gel. We used the DNEasy Plant Mini Kit (Qiagen) for DNA extraction. We homogenised the leaf samples in a dry state, i.e. without adding the buffer prior to homogenisation. We used the same primers and PCR conditions as for the pollen samples described above. The PCR products were sequenced using Sanger sequencing by Macrogen Europe (Netherlands).

We complemented our database by ITS2 sequences from GenBank for those plant species we did not sample in the field. We searched for ITS2 sequences of individual plant species and carefully verified the reliability of records for each species to prevent errors from creeping into our reference database. We aligned the sequences in Geneious using the Geneious Aligner and resolved instances of suspected errors on a case by case basis, particularly by checking the sources of the sequences (data from papers by taxonomists were deemed more reliable than data from ecological surveys, samples from geographically close locations were deemed more relevant, etc.). Public DNA databases are known to contain numerous misidentified records and other types of errors (Bridge et al., 2003). Limiting their impact on our analyses was thus important for confidence in our results.

### Data analysis

We used the Obitools software (Boyer et al., 2016) for bioinformatic processing of the metabarcoding data following general recommendations for filtering and cleaning the sequence data according to De Barba et al. (2014) and Taberlet et al. (2018). We first merged the forward and reverse reads and removed low-quality reads with score<40 or score norm<3.9. We then assigned the reads to samples (keeping the three PCR replicates per sample separate) based on the tag sequences and removed reads shorter than 100 bp, based on available data on the length of the ITS2 region in vascular plants (Chen et al., 2010). We then dereplicated the reads to obtain the list of unique sequences and their abundance in each sample and PCR replicate. We examined results for the blanks and found that the number of reads in blanks ranged from 0 to 4, so we conservatively discarded all sequences with <= 5 reads for each individual sample/PCR replicate to remove sequencing errors caused by tag jumps (De Barba et al., 2014). We then proceeded with sequence identification.

We used our reference database to identify the ITS2 sequences from the samples. We used the *ecotag* function in Obitools to compare each unique ITS2 sequence from the samples with sequences in the reference database. Sequences were identified at the species or genus level with 0.95 as the minimum sequence similarity threshold for taxonomic assignment. To account for possible incompleteness of our reference database, we examined unidentified sequences and attempted to identify them using BLAST search of the GenBank nucleotide database (https://blast.ncbi.nlm.nih.gov/). We found only a few sequences with low read count not matching data from our reference database which could be identified using BLAST. We updated our reference database with these sequences after verifying that the species concerned are known to occur in the wider area around our study site according to botanical records (Grulich, 1997; Wild et al., 2019) or could plausibly occur there. We then reran the sequence identification procedure with the updated reference database. The final outcome of species identification was the number of reads per species (or genus) for each sample and PCR replicate.

The next step was a comparison of the three independent PCR replicates for each sample to identify potentially failed or otherwise unreliable PCR replicates (Taberlet et al., 2018). We calculated the pairwise overlap of the similarity of the plant species composition for all three combinations of the three PCR replicates for each sample using Pianka’s overlap index (Pianka, 1973) calculated using the EcoSimR library (Gotelli et al., 2015) in R 4.0.2 (R Core Team, 2020). The vast majority of comparisons had overlap >0.99 and the smallest value of an overlap between any two PCR replicates of the same sample was 0.94 in samples from the nests and 0.97 in samples from the bodies of foraging females, indicating that different PCR replicates of the same sample gave highly consistent results in all cases. We then averaged the proportions of reads in the three PCR replicates for each sample for the downstream analyses.

Statistical analyses of the data were done in R 4.0.2 (R Core Team, 2020). We used the Pianka’s overlap index (Pianka, 1973) as a measure of similarity in pollen composition among nests, brood cells, and samples from individual foraging trips. To analyse interspecific resource partitioning, we used non-metric multidimensional scaling (NMDS) in two dimensions and ANOSIM - a permutational analysis of similarity (Clarke, 1993), both implemented in the vegan library for R (Oksanen et al., 2019), using Pianka’s overlap index as a measure of similarity in pollen composition among the three *Ceratina* species. For the analysis of pollen from the nests, we averaged the proportion of reads per plant species across all brood cells in each nest and we analogously aggregated repeated samples of pollen from the bodies of individual females. These aggregated data were used in NMDS and ANOSIM. We also used generalised linear models (GLM) to compare the values of Shannon’s H’ index and its components, i.e. the number of plant species and evenness, among nests of the three *Ceratina* species. We included estimated nest age (average age of the brood cells based on the developmental stage) as a covariate to account for possible phenological shifts. We used Gaussian error distribution in the analysis of Shannon’s H’ index and evenness and overdispersed Poisson distribution (quasipoisson) for the number of plant species. We used analogously constructed generalised linear mixed models (GLMM) to analyse these data at the level of individual brood cells, where the nest ID was used as a factor with random effect.

Additional analyses focused on within-individual and between-individual variation in foraging. We used repeatability analysis (Nakagawa and Schielzeth, 2010) to evaluate individual-level differences in specialisation and foraging preferences. Specifically, this analysis compared variation in the Shannon’s H’ index and its components, the number of plant species and evenness, among pollen samples from brood cells from the same nest and among different nests, separately for each of the three *Ceratina* species. We analogously analysed data from pollen samples taken from the bodies of foraging females. To calculate repeatability, we used GLMM fitted by Markov chain Monte Carlo (MCMC), which provides reliable variance estimates, in rptR library for R (Schielzeth and Nakagawa, 2013). We used a GLMM with Gaussian error distribution for Shannon’s H’ index and evenness and Poisson distribution for the number of plant species. We also evaluated individual-level variation in the composition of the pollen samples using partial Mantel tests implemented in the vegan library for R (Oksanen et al., 2019). For pollen data from the nests, we constructed a dissimilarity matrix based on the overlap of the composition of the samples among all combinations of the brood cells across all nests of the same species. Another dissimilarity matrix contained the differences in estimated cell age. We then used a partial Mantel test (Pearson’s correlation, 9999 permutations) to test whether the dissimilarity of pollen composition among brood cells from the same nest differed from the dissimilarity of pollen composition among brood cells from different nests, conditioned on differences in estimated brood cell age. We did this analysis separately for each of the three *Ceratina* species. We did the same type of analysis with data on the composition of pollen samples obtained from the bodies of foraging females.

## RESULTS

Most pollen sequences were identified at the species level, with a few exceptions, e.g. *Rubus* s p. and *Hypericum* sp., where we achieved genus-level identification. Specifically, 90.9% of reads after quality filtering in samples from the nests of *Ceratina* spp. were identified at the species level, while 9.1% of reads were identified at the genus level and a mere 0.03% remained unidentified. In samples from the bodies of *Ceratina* females, 92.2% of reads after quality filtering were identified at the species level, 7.8% at the genus level and 0.01% were unidentified.

### Interspecific resource partitioning

We found clear interspecific differences in pollen composition in nests from the three *Ceratina* species as well as interspecific differences in their level of specialisation. Overall pollen composition in nests of the three *Ceratina* species, expressed as the mean proportion of reads identified as individual plant species, is summarised in Fig. 1. Nests of *C*. *chalybea* and the other two species were separated by a NMDS analysis in two dimensions (Fig. 2), while pollen composition in nests of *C*. *cucurbitina* overlaped with *C*. *nigrolabiata*. Notably, there was also a much higher spread among individual nests in *C*. *chalybea* and *C*. *nigrolabiata* compared to *C*. *cucurbitina*, see Fig. 2, but this could be partly a consequence of a lower number of observations for *C*. *cucurbitina*. Differences in pollen composition from nests of the three *Ceratina* species were strongly supported by ANOSIM, a permutational analysis of similarity, using Pianka’s overlap index as a measure of similarity in pollen composition (*R* = 0.385, *P* < 0.0001, 9999 permutations).

**Figure 1.**
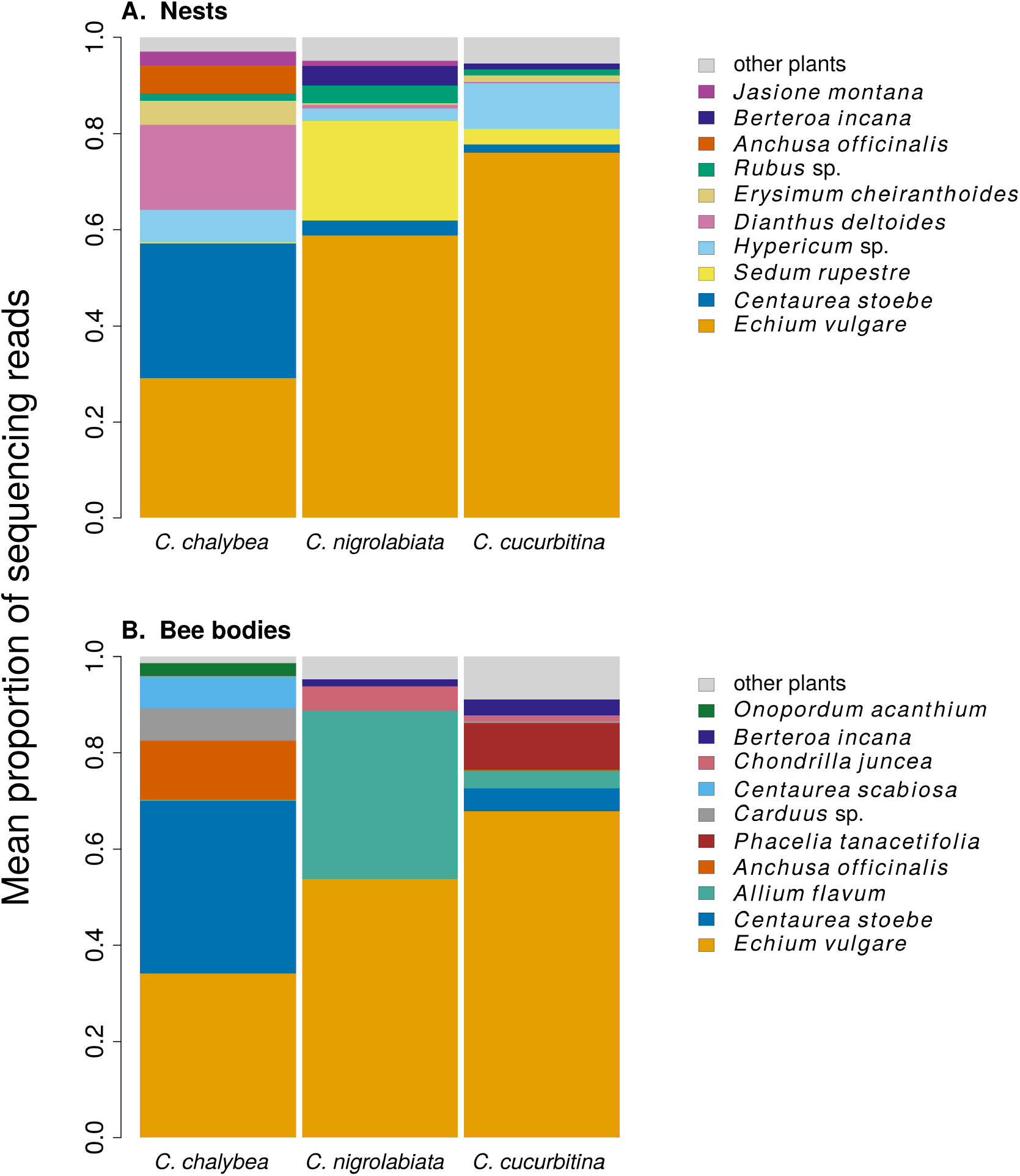
Overall pollen composition of samples from nests and bodies of the three *Ceratina* species. The composition of pollen samples from the nests (A.) and samples collected from the bodies (B.) of females of individual species was calculated as the mean proportion of reads assigned to individual plant species. Plant species are sorted from the bottom up according to their total number of reads in samples from all three *Ceratina* species. Ten species with the highest numbers of reads in samples from the nests (A.) and females bodies (B.) are distinguished by colours, the remaining species are pooled and shown in light grey for clarity.

**Figure 2.**
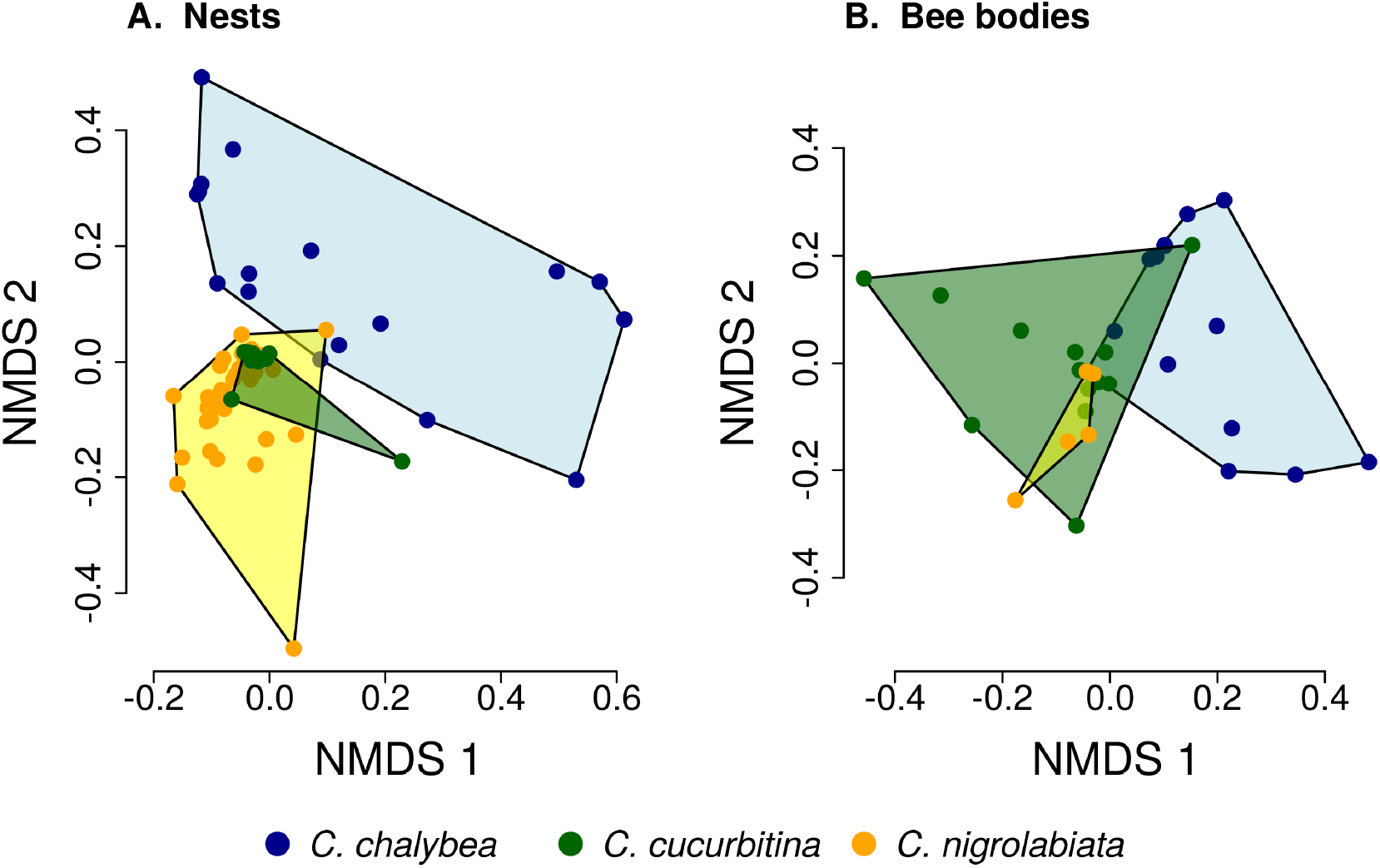
Similarity of pollen composition in individual nests within and among the three *Ceratina* species. Results of Non-metric Multidimensional Scaling (NMDS) showing the similarity of pollen composition of samples from individual nests (A.) and bodies of individual female bees (B.) in two dimensions. The polygons delimit the area containing samples from nests or female bee bodies of individual *Ceratina* species. The position of nests or individual bees is shown by coloured points.

Pollen composition in samples collected four weeks later from the bodies of female bees when returning from a foraging trip to the nest shows patterns consistent with data on pollen composition from the nests (Fig. 1 and Fig. 2). Samples from *C*. *chalybea* were again separated from the other two species by a NMDS analysis (Fig. 2). Differences in pollen composition of samples from bodies of the three *Ceratina* species were also strongly supported by ANOSIM (*R* = 0.197, *P* = 0.002, 9999 permutations).

We observed interspecific differences in pollen diversity ( Fig. 3) measured by the Shannon’s H’ index at the level of entire nests (GLM, *F*_2,63_ = 3.41, *P* = 0.040) and to a limited degree at the level of individual brood cells (GLMM, 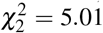, *P* = 0.082). Of the two components of the Shannon’s H’ index, i.e. the number of plant species in a sample and evenness of species composition, only the later differed among the three *Ceratina* species. There was little evidence for interspecific differences in the number of plant species per nest (GLM, *F*_2,63_ = 1.35, *P* = 0.268) or in individual brood cells (GLMM, 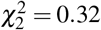, *P* = 0.853). On the other hand, we found clear differences in evenness among the three species at the level of nests (GLM, *F*_2,63_ = 4.54, *P* = 0.015) as well as brood cells (GLMM, 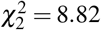, *P* = 0.012), see Fig. 3. Data on pollen diversity in samples collected from the bodies of female *Ceratina*, i.e. pollen collected during a single foraging trip, showed no significant differences among the three species ( Fig. 3).

**Figure 3.**
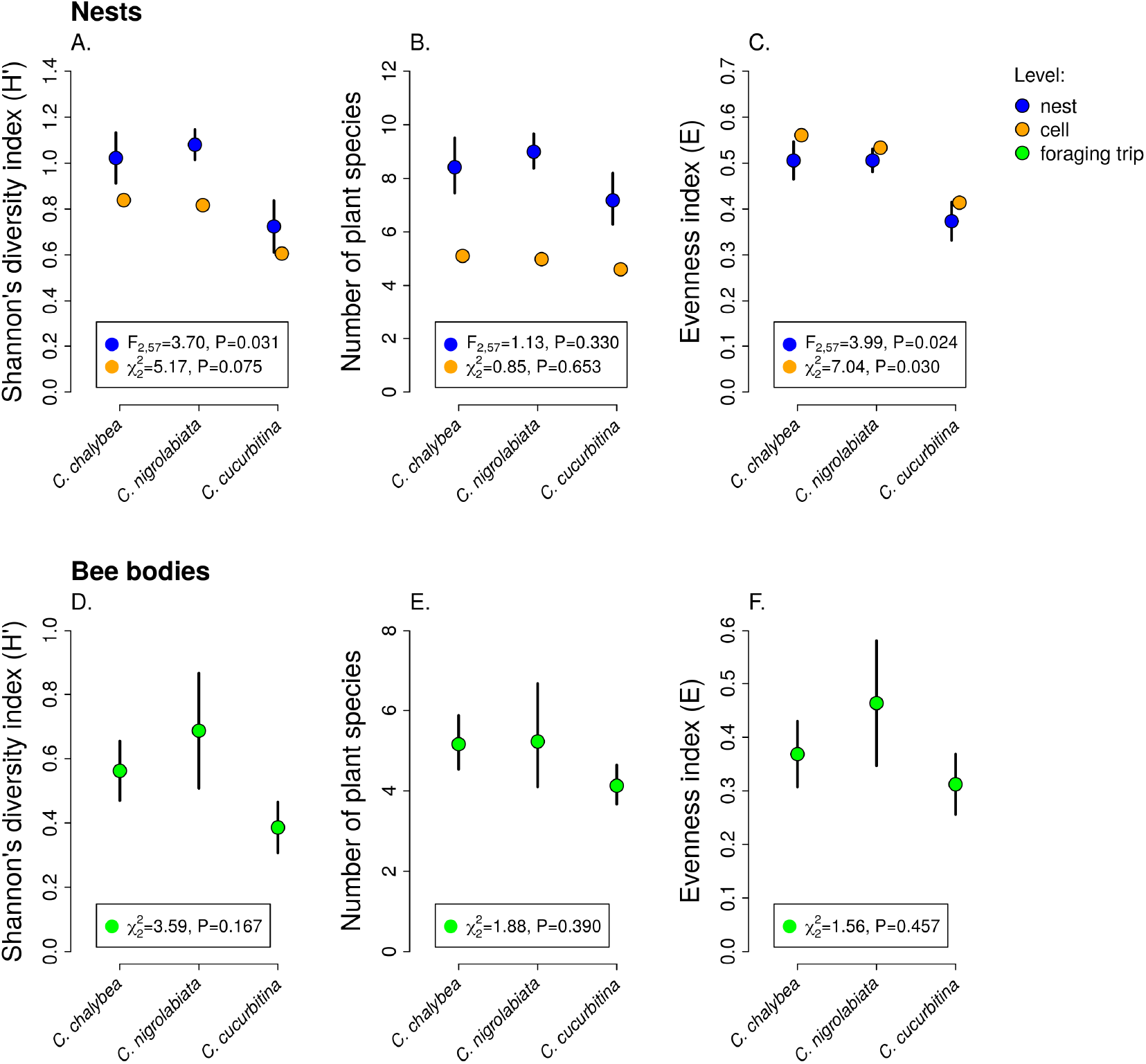
Pollen diversity of samples at the level of nests, brood cells, and individual foraging bouts. Shannon’s diversity index, plant species richness, and evenness (*mean* ± *SE*) calculated from the proportions of reads identified as individual plant species at the level of individual nests and individual brood cells for the three *Ceratina* species (A.-C.) and for individual foraging bouts (D.-F.). The *F* and *P* values refer to the results of GLM and the *χ*^2^ and *P* values refer to the results of GLMM (see Methods and Results).

### Individual-level differences in specialisation and foraging preferences

Females of all three *Ceratina* species showed consistent individual-level differences in their level of specialisation when collecting pollen (Table 1). We found high levels of repeatability of the Shannon’s H index of pollen samples in brood cells from individual nests in all three species (median 0.47-0.70), i.e. brood cells in some nests had consistently higher pollen diversity than brood cells in other nests of the same species. Among the two components of the diversity index (Shannon’s H), evenness was more strongly repeatable than the number of plant species per brood cell, which had high repeatability only in *C*. *chalybea* (Table 1).

**Table 1.** Repeatability analysis shows consistent differences among individuals in measures of foraging specialisation. Results of tests of repeatability of pollen diversity in individual females of the three *Ceratina* species are reported. Mean values of repeatability and 95% credible intervals are reported. Values of repeatability significantly larger than zero (shown in bold) mean that variance of Shannon’s H’, plant species richness, or evenness of pollen composition in brood cells within the same nest (or repeated samples from the body of the same female) was significantly smaller than variance among different nests (or bodies of different females) - this is evidence of consistent among-individual differences. The number of samples from the bodies of *C*. *nigrolabiata* was not sufficient for analysis.

|  | *C. chalybea* | *C. nigrolabiata* | *C. cucurbitina* |
| --- | --- | --- | --- |
| **Nests** |  |  |  |
| Diversity index (Shannon's H) | **0.77** [0.489, 0.878] | **0.45** [0.273, 0.623] | **0.54** [0.280, 0.816] |
| Species richness | **0.53** [0.244, 0.764] | 0.001 [0, 0.171] | 0.28 [0, 0.583] |
| Evenness index (E) | **0.36** [0.204, 0.648] | **0.36** [0.203, 0.517] | **0.37** [0.197, 0.700] |
| **Bee bodies** |  |  |  |
| Diversity index (Shannon's H) | **0.46** [0.118, 0.750] | NA | **0.29** [0.058, 0.597] |
| Species richness | 0.002 [0, 0.509] | NA | 0.004 [0, 0.593] |
| Evenness index (E) | **0.50** [0.191, 0.767] | NA | **0.64** [0.226, 0.835] |

We also found high repeatability of the Shannon’s H and evenness in pollen repeatedly sampled from the bodies of individual females returning from a foraging trip (Table 1). We note that 50% of individuals were recaptured on the same day and 50% on multiple different days in *C*. *chalybea* as well as in *C*. *cucurbitina*. Only a single *C*. *nigrolabiata* was recaptured twice, while other individuals were captured only once, so we had to exclude *C*. *nigrolabiata* from analyses on individual-level differences, because they require repeated sampling of the same individuals.

Individual females of *C*. *chalybea* and *C*. *nigrolabiata*, but not *C*. *cucurbitina*, showed consistent individual-level differences in the composition of pollen contained in brood cells in their nests according to partial Mantel tests conditioned on the temporal distance of the samples (estimated age of brood cells). Similarity in pollen composition between brood cells from the same nest compared to brood cells from different nests was stronger in *C*. *chalybea* (*r* = 0.20, *P* < 0.0001) than in *C*. *nigrolabiata* (*r* = 0.042, *P* < 0.0001), and negligible in *C*. *cucurbitina* (*r* = 0.029, *P* = 0.104), based on 9999 permutations in all cases (Fig. 4). Partial Mantel test accounted for differences in the age of different nests, i.e. changes due to phenology. Indeed, pairs of brood cells with larger differences in their estimated age had more dissimilar pollen composition in all three species (partial Mantel test of the dependence of the dissimilarity of pollen composition on the difference in brood cell age conditioned on whether the pairwise brood cell combination came from the same or different nests, 9999 permutations): *C*. *chalybea* (*r* = 0.234, *P* = 0.0031), *C*. *nigrolabiata* (*r* = 0.151, *P* = 0.0006), and *C*. *cucurbitina* (*r* = 0.424, *P* < 0.0001).

**Figure 4.**
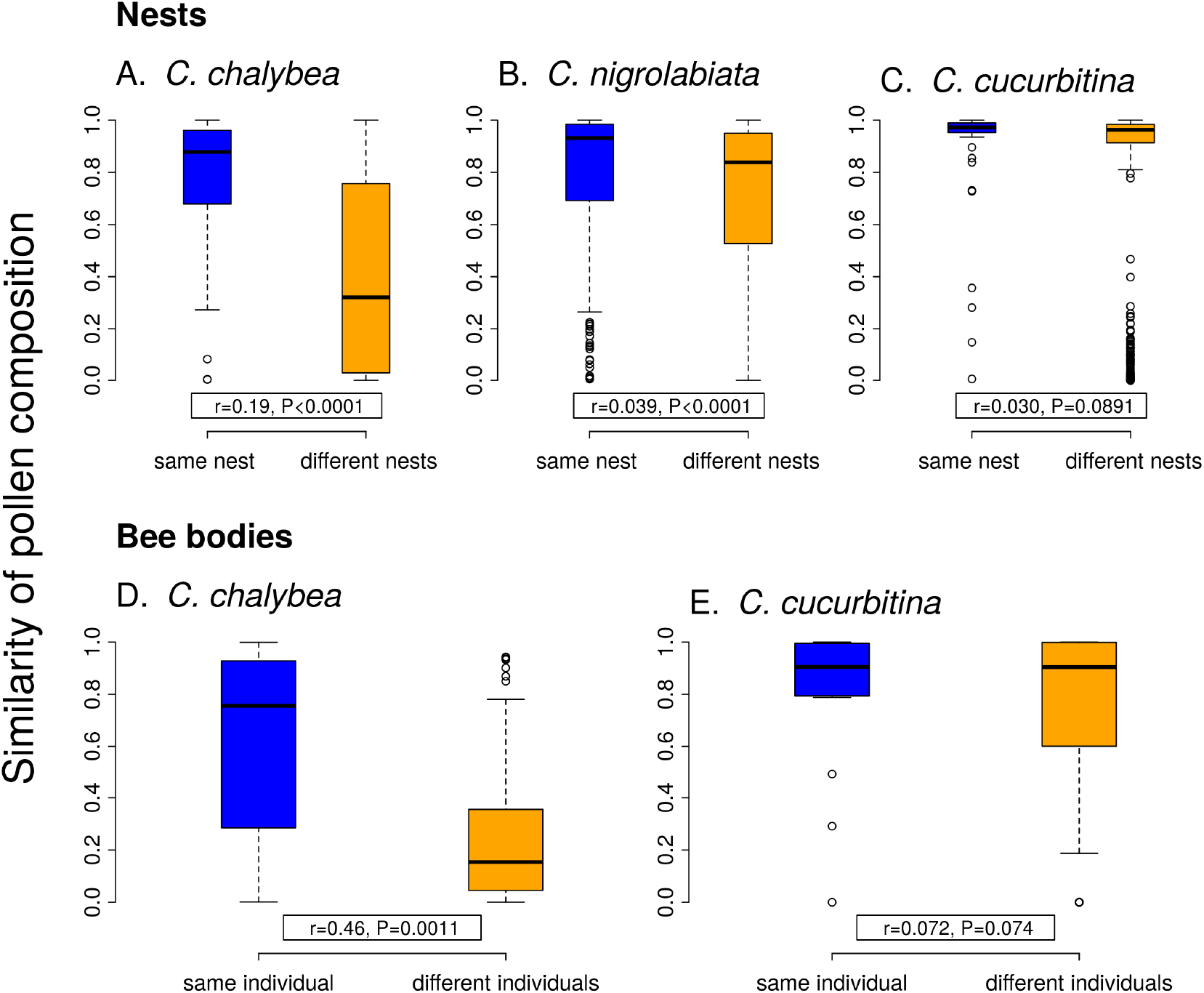
Within-individual and between-individual variation in pollen composition. Higher similarity of pollen composition (Pianka’s overlap index) between brood cells from the same nest compared to brood cells from different nests (A.-C.) indicates consistent differences in foraging preferences among individual females. Similarity in pollen composition is analogously compared between repeated samples from the bodies of foraging females (D.-E.). The number of samples from bodies of *C*. *nigrolabiata* was not sufficient for analysis. Median and interquartile range is shown in the boxplots. The *r* and *P* values refer to the results of partial Mantel tests (see Methods and Results).

Similarly to data from the nests, we found consistent individual-level differences in the composition of pollen sampled directly from bodies of repeatedly captured females of *C*. *chalybea* returning from a foraging trip (partial Mantel test, *r* = 0.460, *P* = 0.0006, 9999 permutations), but not in *C*. *cucurbitina* (*r* = 0.071, *P* = 0.084). There was no effect of temporal distance (the number of days between collecting the samples) on similarity of pollen composition in both *C*. *chalybea* (*r* = −0.236, *P* = 0.999) and *C. cucurbitina* (*r* = −0.006, *P* = 0.446) (partial Mantel test conditioned on whether pairwise sample combination came from the same or different individual, 9999 permutations). That means that similarity in the composition of pollen collected from the same individual did not depend on whether the individual was recaptured the same day or several days apart.

## DISCUSSION

### Individual-level consistency and among-individual variation in specialisation and foraging preferences

In this study, we tested how diet breadth and selectivity of three co-occurring species of bees foraging for pollen varies across various levels of aggregation from a single foraging bout, through an individual’s lifetime, to the population and species level. We found that the females of all three species displayed consistent among-individual differences in foraging specialisation at the short temporal scale of individual foraging bouts as well as at a longer temporal scale represented by pollen provisions accumulated in brood cells in their nests. Moreover, larger among-individual differences at the intraspecific level translated into lower specialisation at the species level. Individuals of the more generalised species (*C*. *chalybea* and to a lesser extent *C*. *nigrolabiata*) displayed significant among-individual differences in foraging preferences and had larger within-individual variation. On the other hand, individuals of the most specialised species (*C*. *cucurbitina*) were extremely consistent in their foraging preferences both at the within-individual and among-individual level. Differences in foraging strategies of the three species imply that they may play different functional roles in the local plant-pollinator network.

Our data thus support the conceptual scheme of varying levels of specialisation at different levels of aggregation presented by Brosi (2016) based on earlier studies on individual-level variation of diet breadth in various consumers (Bolnick et al., 2003; Araújo et al., 2011). As predicted, we found that the three species of solitary bees of the genus *Ceratina* were more specialised at the level of individual foraging bouts than over longer time scales, based on a comparison of pollen diversity in samples from single foraging trips, pollen provisions in individual brood cells accumulated over a few days, and pollen aggregated in entire nests collected by a single female over the period of many days, see also Kobayashi-Kidokoro and Higashi (2010). Moreover, we found consistent among-individual differences in their specialisation and foraging preferences. Hence, some individuals were consistently more specialised than other individuals of the same species (repeatability analysis; Table 1) and they foraged on a different set of plant species (Fig. 4). Our study thus provides unambiguous evidence of consistent individual-level specialisation in pollinators following early observations by (Heinrich, 1976) in bumblebees and recent evidence in butterflies (Szigeti et al., 2019).

At the species level, we found a clear distinction in the level of specialisation and foraging preferences among the three *Ceratina* species. *C*. *chalybea* and *C*. *nigrolabiata* were more generalised than *C*. *cucurbitina*. These differences stem mostly from varying strength of among-individual differences at the intraspecific level (Fig. 4). While we observed large differences in pollen composition among different individuals of *C*. *chalybea*, among-individual differences were smaller in *C*. *nigrolabiata* and virtually absent in *C*. *cucurbitina*, where all individuals were extremely consistent in their specialisation and foraging preferences. Hence, we detected large differences in specialisation among the three species at the aggregated species level despite the relatively small and statistically insignificant differences among the three species in specialisation at the level of individual foraging bouts. Higher generalisation at the species level thus stemmed from larger among-iúndividual variation in diets as observed in other types of consumers, such as predators (Bolnick et al., 2003; Araújo et al., 2011).

Interestingly, *C*. *nigrolabiata* has a longer duration of foraging trips compared to the other two species, likely because it has biparental care and the male guards the nest while the female is foraging (Mikát et al., 2019). Lower time constraints on foraging could promote specialisation on the most rewarding resources (Lucas, 1983), but we observed a similar level of specialisation in the three species during single foraging bouts. This may be driven by the same balance of energetic costs and benefits of selective feeding in all three species (Emlen, 1966; Grüter and Ratnieks, 2011). In contrast, two closely related honeybee species, *Apis cerana* and *A. mellifera*, differ in their levels of flower constancy according to Wells and Rathore (1994). However, we detected differences in individual-level consistency over a longer time scale among the three *Ceratina* species, despite the fact that they were studied at the same site and exposed to the same abundance and composition of resources. These differences may stem from different nutritional demands of the larvae or different levels of foraging flexibility in the three species (Grüter and Ratnieks, 2011).

We also conclude that there was a certain level of resource partitioning among the three *Ceratina* species. While pollen from e.g. *Echium vulgare* was found in pollen provisions of all three species, many plant species were found exclusively in pollen provisions of only one or two of the three *Ceratina* species. For example, pollen of *Centaurea stoebe* and *Dianthus deltoides* was common in the samples from *C*. *chalybea* but almost absent in samples from the other two *Ceratina* species, where it was replaced by pollen of *Sedum rupestre*, *Alium flavum*, and other plants. Such pattern of resource partitioning could be caused by different preferences for floral traits (Junker et al., 2013; Klecka et al., 2018a), variation in preferred plant height (Klecka et al., 2018b), or by interspecific competition (Schoener, 1974; Palmer et al., 2003), as demonstrated previously in bumblebees (Inouye, 1978; Graham and Jones, 1996).

An interesting fact is that pollen of *Echium vulgare* was the dominant source of pollen for all three *Ceratina* species, based on the proportion of sequencing reads. The pollen of *E. vulgare* has a very high protein content (35% crude protein in the dry matter according to Somerville and Nicol (2006)), which makes it a potentially excellent resource for bees, but it contains high concentrations of pyrrolizidine alkaloids (Boppré et al., 2008; Lucchetti et al., 2016; Trunz et al., 2020) toxic to insects (Narberhaus et al., 2005; Macel, 2011). Only a restricted range of solitary bee species can successfully develop on the pollen of *E. vulgare* (Praz et al., 2008; Sedivy et al., 2011; Trunz et al., 2020). In particular, some species of the genus *Hoplitis* (Hymenoptera: Megachilidae) are specialised on *Echium* and other plants in the family Boraginaceae (Sedivy et al., 2013), which are also known to contain pyrrolizidine alkaloids (El-Shazly et al., 1998; El-Shazly and Wink, 2014; Trunz et al., 2020). Our results suggest that the three species of *Ceratina* we studied also have physiological adaptations to develop on the pollen of *E. vulgare*, which allows them to utilise its protein-rich pollen Somerville and Nicol (2006).

### Implications of variation in specialisation and foraging preferences across individuals and temporal scales

Foraging behaviour of flower visitors has important consequences for reproduction of entomophilous plants. From the plant’s perspective, high level of specialisation of it’s pollinators intuitively seems desirable. Specialised pollinators may be more effective than generalists, i.e. they provide higher single visit contribution to plant reproductive fitness (Larsson, 2005; McIntosh, 2005), although e.g. specialised solitary bees often remove more pollen per flower visit than generalists, which increases the costs for the plants (Larsson, 2005; Parker et al., 2016). However, it is important to emphasise that specialisation specifically at the level of individual foraging bouts, i.e. high flower constancy, matters for pollination because it ensures that pollen is transferred between flowers of the same plant species (Brosi, 2016) and it minimises heterospecific pollen transfer which may decrease both the male fitness of the donor plant and the female fitness of the recipient plant (Waser, 1978; Morales and Traveset, 2008). Hence, even a flower visitor which is generalised at a longer temporal scale (e.g. during it’s lifetime), may be a highly efficient pollinator if it temporarily specialises on a single plant species during a foraging bout (Brosi, 2016; Szigeti et al., 2019). This suggests that pollination efficiency of the three *Ceratina* species we studied may be similar despite their large differences in specialisation at the species level, because they had a comparably high level of specialisation during individual foraging bouts.

Despite the varying level of specialisation, individual brood cells always contained a mixture of pollen of several plant species, which is in line with data on other *Ceratina* species (Kobayashi-Kidokoro and Higashi, 2010; Lawson et al., 2016; McFrederick and Rehan, 2016). The effect of the composition of pollen provisions on the larval development and survival is not straightforward (Nicholls and Hempel de Ibarra, 2017). It has been demonstrated that higher protein content provides benefits for larval development with positive effects persisting to adulthood (Roulston and Cane, 2002; Li et al., 2012). Accordingly, the most utilised plant by *Ceratina* in our study was *Echium vulgare*, whose pollen is one of the most protein-rich among all plants Roulston and Cane (2000); Somerville and Nicol (2006). However, pollen from different plants varies widely not only in protein content, but also in energetic value, lipid contents, etc. (Roulston and Cane, 2000; Somerville and Nicol, 2006; Brodschneider and Crailsheim, 2010; Vaudo et al., 2020). A mixed diet thus may be beneficial for larval development (Eckhardt et al., 2014), because it could either better satisfy their nutritional needs or dilute toxins present in some of the food sources (Lefcheck et al., 2013; Eckhardt et al., 2014; Vaudo et al., 2016). Although we still know little about the importance of resource diversity for the nutrition of solitary bees, it seems that restricted plant diversity caused by climate change or land use change may have a detrimental effect not only on specialised but also on generalised pollinators by affecting their nutrition (Vaudo et al., 2016, 2020), which should be recognised in planning conservation actions (Vaudo et al., 2015).

Variation in pollen composition among nests built by different females and even among brood cells in the same nest could lead to differences in the growth and traits of the developing larvae. At the intraspecific level, variation in pollen provisions among nests is an example of maternal effects (Bernardo, 1996): the development and traits of the offspring may be driven by individual foraging preferences of their mother. This way, among-individual variation in foraging preferences may promote phenotypic plasticity in the next generation, which may affect evolutionary changes in the solitary bees (Räsänen and Kruuk, 2007). At an even finer level, variation in pollen composition among brood cells in the same nest could play an important role in the evolution of eusociality in bees - maternal manipulation of the provisions is known to affect the development of the offspring leading to the production of workers in a facultatively eusocial bee *Megalopta genalis* (Halictidae) (Kapheim et al., 2011) and to the production of a dwarf eldest daughter which acts as a worker in *Ceratina calcarata* in North America (Lawson et al., 2016). However, there is no evidence of such maternal manipulation in the three species we studied (Mikát et al., 2020a,b). It is also possible that variation in the composition of the pollen provisions may affect the development of the larvae not only directly by differences in nutritional value, but also indirectly by differences in the composition of bacterial communities in the pollen provisions (McFrederick and Rehan, 2016). We are only beginning to understand such implications of individual foraging behaviour, so there is a number avenues for future research.

### Utility and caveats of the DNA metabarcoding approach

Obtaining such detailed insights was facilitated by the use of a rigorous DNA metabarcoding protocol with different types of controls and by a creation of a local reference database which allowed us to identify pollen DNA sequences with high level of precision (Zinger et al., 2019). In our case, >90% of reads were identified at the species level and almost all the remaining reads at the genus level. This level of precision is unusual when using ITS2 as a marker for plant identification, because many closely related plant species cannot be confidently distinguished. Detailed knowledge of the local flora is thus an important prerequisite for pollen DNA metabarcoding studies where detailed species level data are needed (Biella et al., 2019b). We could rely on a long tradition of botanical surveys at the study site and its surroundings (Grulich, 1997) to obtain an exhaustive list of plant species known from the area. However, compiling a database of ITS2 sequences was still complicated by the high frequency of erroneous or spurious records in public databases.

A caveat of using DNA metabarcoding to analyse the composition of pollen samples is that it is not entirely quantitative, i.e. the proportion of reads belonging to a plant species is generally not a good proxy for the pollen mass or the number of pollen grains because of different DNA contents per unit mass, amplification bias, etc. (Bell et al., 2016). However, the number of pollen grains of individual species per sample correlates positively with the number of sequencing reads (Keller et al., 2015), which suggests that the proportion of reads may provide at least semi-quantitative information. Importantly, this uncertainty is problematic for absolute quantification, i.e., we cannot make conclusions about the amount of pollen collected by the bees based on the number of reads, but it does not invalidate relative comparisons among samples, which is what we focused on in our analyses.

## CONCLUSIONS

In conclusion, we showed that three species of solitary bees of the genus *Ceratina* were more specialised at the level of individual foraging bouts than over longer time scales. Moreover, we found consistent among-individual differences in their specialisation and foraging preferences. Hence, some individuals were consistently more specialised than other individuals of the same species and collected pollen from a different set of plant species. Our study thus provides evidence of consistent individual-level specialisation in pollinators. Moreover, higher generalisation at the species level stemmed from larger among-individual variation in diets as observed in other types of consumers, particularly predators. More detailed knowledge of specialisation and foraging preferences of pollinators across different spatial and temporal scales, from an individual foraging bout to the species level, is necessary to understand plant-flower visitor networks from the functional perspective (Brosi, 2016) and to forecast the consequences of various environmental changes on the robustness of plant-pollinator networks which is mediated by foraging flexibility of pollinators (Biella et al., 2019a, 2020).

## ACKNOWLEDGEMENTS

We would like to thank Tereza Hadravová, Jitka Waldhauserová, Marcela Dokulilová, Kateřina Čermáková, Daniela Reiterová, Šimon Zeman, Vojtěch Brož, and Celie Korittová for their help in the field. JK also thanks Pierre Taberlet, Eric Coissac, and their colleagues for sharing their invaluable expertise during the seventh DNA metabarcoding Spring School at Porto, Portugal, in May 2017. The study was supported by the Czech Science Foundation (projects GJ17-24795Y and GA20-14872S).

## DATA AVAILABILITY

Raw data are available in Figshare at https://www.doi.org/10.6084/m9.figshare.13850324. DNA sequences of plants generated during this project are available in BOLD and Genbank; their list is included in Figshare at https://www.doi.org/10.6084/m9.figshare.13850324.

## SUPPLEMENTAL INFORMATION

Supplemental information is available in Figshare at https://www.doi.org/10.6084/m9.figshare.13850324.

## REFERENCES

Amaya-Márquez, M. (2009). Floral constancy in bees: a revision of theories and a comparison with other pollinators. Revista Colombiana de Entomología, 35(2):206–216.

Araújo, M. S., Bolnick, D. I., and Layman, C. A. (2011). The ecological causes of individual specialisation. Ecology Letters, 14(9):948–958.

Bell, K. L., De Vere, N., Keller, A., Richardson, R. T., Gous, A., Burgess, K. S., and Brosi, B. J. (2016). Pollen DNA barcoding: current applications and future prospects. Genome, 59(9):629–640.

Bell, K. L., Fowler, J., Burgess, K. S., Dobbs, E. K., Gruenewald, D., Lawley, B., Morozumi, C., and Brosi, B. J. (2017). Applying pollen dna metabarcoding to the study of plant–pollinator interactions. Applications in Plant Sciences, 5(6):1600124.

Bernardo, J. (1996). Maternal effects in animal ecology. American Zoologist, 36(2):83–105.

Biella, P., Akter, A., Ollerton, J., Nielsen, A., and Klecka, J. (2020). An empirical attack tolerance test alters the structure and species richness of plant-pollinator networks. Functional Ecology, 34(11):2246–2258.

Biella, P., Akter, A., Ollerton, J., Tarrant, S., Janeček, Š., Jersáková, J., and Klecka, J. (2019a). Experimental loss of generalist plants reveals alterations in plant-pollinator interactions and a constrained flexibility of foraging. Scientific Reports, 9:7376.

Biella, P., Tommasi, N., Akter, A., Guzzetti, L., Klecka, J., Sandionigi, A., Labra, M., and Galimberti, A. (2019b). Foraging strategies are maintained despite workforce reduction: A multidisciplinary survey on the pollen collected by a social pollinator. PLoS ONE, 14(11):e0224037.

Bolnick, D. I., Amarasekare, P., Araújo, M. S., Bürger, R., Levine, J. M., Novak, M., Rudolf, V. H., Schreiber, S. J., Urban, M. C., and Vasseur, D. A. (2011). Why intraspecific trait variation matters in community ecology. Trends in Ecology & Evolution, 26(4):183–192.

Bolnick, D. I., Svanbäck, R., Araújo, M. S., and Persson, L. (2007). Comparative support for the niche variation hypothesis that more generalized populations also are more heterogeneous. Proceedings of the National Academy of Sciences, 104(24):10075–10079.

Bolnick, D. I., Svanbäck, R., Fordyce, J. A., Yang, L. H., Davis, J. M., Hulsey, C. D., and Forister, M. L. (2003). The ecology of individuals: incidence and implications of individual specialization. American Naturalist, 161(1):1–28.

Bolnick, D. I., Yang, L. H., Fordyce, J. A., Davis, J. M., and Svanbäck, R. (2002). Measuring individual level resource specialization. Ecology, 83(10):2936–2941.

Boppré, M., Colegate, S. M., Edgar, J. A., and Fischer, O. W. (2008). Hepatotoxic pyrrolizidine alkaloids in pollen and drying-related implications for commercial processing of bee pollen. Journal of Agricultural and Food Chemistry, 56(14):5662–5672.

Boyer, F., Mercier, C., Bonin, A., Le Bras, Y., Taberlet, P., and Coissac, E. (2016). OBITOOLS: A UNIX-inspired software package for DNA metabarcoding. Molecular Ecology Resources, 16(1):176–182.

Bridge, P. D., Roberts, P. J., Spooner, B. M., and Panchal, G. (2003). On the unreliability of published DNA sequences. New Phytologist, 160(1):43–48.

Brodschneider, R. and Crailsheim, K. (2010). Nutrition and health in honey bees. Apidologie, 41(3):278–294.

Brosi, B. J. (2016). Pollinator specialization: from the individual to the community. New Phytologist, 210(4):1190–1194.

Brosi, B. J. and Briggs, H. M. (2013). Single pollinator species losses reduce floral fidelity and plant reproductive function. Proceedings of the National Academy of Sciences, 110(32):13044–13048.

Chen, S., Yao, H., Han, J., Liu, C., Song, J., Shi, L., Zhu, Y., Ma, X., Gao, T., Pang, X., et al. (2010). Validation of the ITS2 region as a novel DNA barcode for identifying medicinal plant species. PLoS ONE, 5(1):e8613.

Clarke, K. R. (1993). Non-parametric multivariate analyses of changes in community structure. Australian Journal of Ecology, 18(1):117–143.

De Barba, M., Miquel, C., Boyer, F., Mercier, C., Rioux, D., Coissac, E., and Taberlet, P. (2014). DNA metabarcoding multiplexing and validation of data accuracy for diet assessment: application to omnivorous diet. Molecular Ecology Resources, 14(2):306–323.

Eckhardt, M., Haider, M., Dorn, S., and Müller, A. (2014). Pollen mixing in pollen generalist solitary bees: a possible strategy to complement or mitigate unfavourable pollen properties? Journal of Animal Ecology, 83(3):588–597.

El-Shazly, A., El-Domiaty, M., Witte, L., and Wink, M. (1998). Pyrrolizidine alkaloids in members of the Boraginaceae from Sinai (Egypt). Biochemical Systematics and Ecology, 26(6):619–636.

El-Shazly, A. and Wink, M. (2014). Diversity of pyrrolizidine alkaloids in the Boraginaceae structures, distribution, and biological properties. Diversity, 6(2):188–282.

Emlen, J. M. (1966). The role of time and energy in food preference. American Naturalist, 100(916):611–617.

Fontaine, C., Collin, C. L., and Dajoz, I. (2008). Generalist foraging of pollinators: diet expansion at high density. Journal of Ecology, 96(5):1002–1010.

Forsman, A. and Wennersten, L. (2016). Inter-individual variation promotes ecological success of populations and species: Evidence from experimental and comparative studies. Ecography, 39(7):630–648.

Gegear, R. and Laverty, T. (1998). How many flower types can bumble bees work at the same time? Canadian Journal of Zoology, 76(7):1358–1365.

Gotelli, N. J., Hart, E. M., and Ellison, A. M. (2015). EcoSimR: Null model analysis for ecological data. R package version 0.1.0.

Goulson, D. and Wright, N. P. (1998). Flower constancy in the hoverflies *Episyrphus balteatus* (Degeer) and *Syrphus ribesii* (L.)(Syrphidae). Behavioral Ecology, 9(3):213–219.

Graham, L. and Jones, K. N. (1996). Resource partitioning and per-flower foraging efficiency in two bumble bee species. American Midland Naturalist, 136(2):401–406.

Groom, S. V. and Rehan, S. M. (2018). Climate-mediated behavioural variability in facultatively social bees. Biological Journal of the Linnean Society, 125(1):165–170.

Grulich, V. (1997). Atlas rozšíření cévnatých rostlin Národního parku Podyjí. Masaryk University, Brno, Czech Republic.

Grüter, C. and Ratnieks, F. L. (2011). Flower constancy in insect pollinators: Adaptive foraging behaviour or cognitive limitation? Communicative & Integrative Biology, 4(6):633–636.

Heinrich, B. (1976). The foraging specializations of individual bumblebees. Ecological Monographs, 46(2):105–128.

Hofstede, F. E. and Sommeijer, M. J. (2006). Effect of food availability on individual foraging specialisation in the stingless bee *Plebeia tobagoensis* (Hymenoptera, Meliponini). Apidologie, 37(3):387–397.

Inouye, D. W. (1978). Resource partitioning in bumblebees: experimental studies of foraging behavior. Ecology, 59(4):672–678.

Junker, R. R., Blüthgen, N., Brehm, T., Binkenstein, J., Paulus, J., Schaefer, M. H., and Stang, M. (2013). Specialization on traits as basis for the niche-breadth of flower visitors and as structuring mechanism of ecological networks. Functional Ecology, 27(2):329–341.

Kapheim, K. M., Bernal, S. P., Smith, A. R., Nonacs, P., and Wcislo, W. T. (2011). Support for maternal manipulation of developmental nutrition in a facultatively eusocial bee, *Megalopta genalis* (Halictidae). Behavioral Ecology and Sociobiology, 65(6):1179–1190.

Keller, A., Danner, N., Grimmer, G., Ankenbrand, M v. d.,, Von Der Ohe, K., Von Der Ohe, W., Rost, S., Härtel, S., and Steffan-Dewenter, I. (2015). Evaluating multiplexed next-generation sequencing as a method in palynology for mixed pollen samples. Plant Biology, 17(2):558–566.

Kitson, J. J., Hahn, C., Sands, R. J., Straw, N. A., Evans, D. M., and Lunt, D. H. (2019). Detecting host–parasitoid interactions in an invasive lepidopteran using nested tagging DNA metabarcoding. Molecular Ecology, 28(2):471–483.

Klecka, J., Hadrava, J., Biella, P., and Akter, A. (2018a). Flower visitation by hoverflies (Diptera: Syrphidae) in a temperate plant-pollinator network. PeerJ, 6:e6025.

Klecka, J., Hadrava, J., and Koloušková, P. (2018b). Vertical stratification of plant–pollinator interactions in a temperate grassland. PeerJ, 6:e4998.

Kobayashi-Kidokoro, M. and Higashi, S. (2010). Flower constancy in the generalist pollinator *Ceratina flavipes* (Hymenoptera: Apidae): an evaluation by pollen analysis. Psyche, 2010:891906.

Larsson, M. (2005). Higher pollinator effectiveness by specialist than generalist flower-visitors of unspecialized *Knautia arvensis* (Dipsacaceae). Oecologia, 146(3):394–403.

Lawson, S. P., Ciaccio, K. N., and Rehan, S. M. (2016). Maternal manipulation of pollen provisions affects worker production in a small carpenter bee. Behavioral Ecology and Sociobiology, 70(11):1891–1900.

Lefcheck, J. S., Whalen, M. A., Davenport, T. M., Stone, J. P., and Duffy, J. E. (2013). Physiological effects of diet mixing on consumer fitness: a meta-analysis. Ecology, 94(3):565–572.

Lewis, A. C. (1986). Memory constraints and flower choice in *Pieris rapae*. Science, 232(4752):863–865.

Li, C., Xu, B., Wang, Y., Feng, Q., and Yang, W. (2012). Effects of dietary crude protein levels on development, antioxidant status, and total midgut protease activity of honey bee (*Apis mellifera ligustica*). Apidologie, 43(5):576–586.

Lucas, J. R. (1983). The role of foraging time constraints and variable prey encounter in optimal diet choice. American Naturalist, 122(2):191–209.

Lucchetti, M. A., Glauser, G., Kilchenmann, V., Dübecke, A., Beckh, G., Praz, C., and Kast, C. (2016). Pyrrolizidine alkaloids from *Echium vulgare* in honey originate primarily from floral nectar. Journal of Agricultural and Food Chemistry, 64(25):5267–5273.

Macel, M. (2011). Attract and deter: a dual role for pyrrolizidine alkaloids in plant–insect interactions. Phytochemistry Reviews, 10(1):75–82.

McFrederick, Q. S. and Rehan, S. M. (2016). Characterization of pollen and bacterial community composition in brood provisions of a small carpenter bee. Molecular Ecology, 25(10):2302–2311.

McIntosh, M. (2005). Pollination of two species of *Ferocactus*: interactions between cactus-specialist bees and their host plants. Functional Ecology, 19(4):727–734.

Mikát, M., Benda, D., Korittová, C., Mrozková, J., Reiterová, D., Waldhauserová, J., Brož, V., and Straka, J. (2020a). Natural history and maternal investment of *Ceratina cucurbitina*, the most common european small carpenter bee, in different european regions. Journal of Apicultural Research.

Mikát, M., Černá, K., and Straka, J. (2016). Major benefits of guarding behavior in subsocial bees: implications for social evolution. Ecology and Evolution, 6(19):6784–6797.

Mikát, M., Franchino, C., and Rehan, S. M. (2017). Sociodemographic variation in foraging behavior and the adaptive significance of worker production in the facultatively social small carpenter bee, *Ceratina calcarata*. Behavioral Ecology and Sociobiology, 71(9):135.

Mikát, M., Janošík, L., Černá, K., Matoušková, E., Hadrava, J., Bureš, V., and Straka, J. (2019). Polyandrous bee provides extended offspring care biparentally as an alternative to monandry based eusociality. Proceedings of the National Academy of Sciences, 116(13):6238–6243.

Mikát, M., Waldhauserová, J., Fraňková, T., Čermáková, K., Brož, V., Zeman, Š., Dokulilová, M., and Straka, J. (2020b). Only mothers feed mature offspring in european *Ceratina* bees. Insect Science.

Morales, C. L. and Traveset, A. (2008). Interspecific pollen transfer: magnitude, prevalence and consequences for plant fitness. Critical Reviews in Plant Sciences, 27(4):221–238.

Nakagawa, S. and Schielzeth, H. (2010). Repeatability for Gaussian and non-Gaussian data: a practical guide for biologists. Biological Reviews, 85(4):935–956.

Narberhaus, I., Zintgraf, V., and Dobler, S. (2005). Pyrrolizidine alkaloids on three trophic levels–evidence for toxic and deterrent effects on phytophages and predators. Chemoecology, 15(2):121–125.

Nicholls, E. and Hempel de Ibarra, N. (2017). Assessment of pollen rewards by foraging bees. Functional Ecology, 31(1):76–87.

Noreika, N., Bartomeus, I., Winsa, M., Bommarco, R., and Öckinger, E. (2019). Pollinator foraging flexibility mediates rapid plant-pollinator network restoration in semi-natural grasslands. Scientific Reports, 9:15473.

Oksanen, J., Blanchet, F. G., Friendly, M., Kindt, R., Legendre, P., McGlinn, D., Minchin, P. R., O’Hara, R. B., Simpson, G. L., Solymos, P., Stevens, M. H. H., Szoecs, E., and Wagner, H. (2019). vegan: Community Ecology Package. R package version 2.5-6.

Palmer, T. M., Stanton, M. L., and Young, T. P. (2003). Competition and coexistence: exploring mechanisms that restrict and maintain diversity within mutualist guilds. American Naturalist, 162(S4):S63–S79.

Parker, A. J., Williams, N. M., and Thomson, J. D. (2016). Specialist pollinators deplete pollen in the spring ephemeral wildflower *Claytonia virginica*. Ecology and Evolution, 6(15):5169–5177.

Pianka, E. R. (1973). The structure of lizard communities. Annual Review of Ecology and Systematics, 4(1):53–74.

Praz, C. J., Müller, A., and Dorn, S. (2008). Specialized bees fail to develop on non-host pollen: do plants chemically protect their pollen? Ecology, 89(3):795–804.

R Core Team (2020). R: A Language and Environment for Statistical Computing. R Foundation for Statistical Computing, Vienna, Austria.

Räsänen, K. and Kruuk, L. (2007). Maternal effects and evolution at ecological time-scales. Functional Ecology, 21(3):408–421.

Rehan, S. M., Leys, R., and Schwarz, M. P. (2012). A mid-cretaceous origin of sociality in xylocopine bees with only two origins of true worker castes indicates severe barriers to eusociality. PLoS ONE, 7(4).

Rehan, S. M. and Richards, M. H. (2010). Nesting biology and subsociality in *Ceratina calcarata* (Hymenoptera: Apidae). The Canadian Entomologist, 142(1):65–74.

Roughgarden, J. (1972). Evolution of niche width. American Naturalist, 106(952):683–718.

Roughgarden, J. (1974). Niche width: biogeographic patterns among anolis lizard populations. American Naturalist, 108(962):429–442.

Roulston, T. H. and Cane, J. H. (2000). Pollen nutritional content and digestibility for animals. In Dafni, A., Hesse, M., and Pacini, E., editors, Pollen and Pollination, pages 187–209. Springer, Vienna, Austria.

Roulston, T. H. and Cane, J. H. (2002). The effect of pollen protein concentration on body size in the sweat bee *Lasioglossum zephyrum* (Hymenoptera: Apiformes). Evolutionary Ecology, 16(1):49–65.

Schielzeth, H. and Nakagawa, S. (2013). rptR: Repeatability for Gaussian and non-Gaussian data. R package version 0.6.405/r52.

Schoener, T. W. (1974). Resource partitioning in ecological communities. Science, 185(4145):27–39.

Sedivy, C., Dorn, S., Widmer, A., and Müller, A. (2013). Host range evolution in a selected group of osmiine bees (Hymenoptera: Megachilidae): the Boraginaceae-Fabaceae paradox. Biological Journal of the Linnean Society, 108(1):35–54.

Sedivy, C., Müller, A., and Dorn, S. (2011). Closely related pollen generalist bees differ in their ability to develop on the same pollen diet: evidence for physiological adaptations to digest pollen. Functional Ecology, 25(3):718–725.

Sickel, W., Ankenbrand, M. J., Grimmer, G., Holzschuh, A., Härtel, S., Lanzen, J., Steffan-Dewenter, I., and Keller, A. (2015). Increased efficiency in identifying mixed pollen samples by meta-barcoding with a dual-indexing approach. BMC Ecology, 15(1):20.

Slaa, J. E., Cevaal, A., and Sommeijer, M. J. (1998). Floral constancy in *Trigona* stingless bees foraging on artificial flower patches: a comparative study. Journal of Apicultural Research, 37(3):191–198.

Somerville, D. and Nicol, H. (2006). Crude protein and amino acid composition of honey bee-collected pollen pellets from south-east Australia and a note on laboratory disparity. Australian Journal of Experimental Agriculture, 46(1):141–149.

Strickler, K. (1979). Specialization and foraging efficiency of solitary bees. Ecology, 60(5):998–1009.

Szigeti, V., Kőrösi, Á., Harnos, A., and Kis, J. (2019). Lifelong foraging and individual specialisation are influenced by temporal changes of resource availability. Oikos, 128(5):649–658.

Taberlet, P., Bonin, A., Coissac, E., and Zinger, L. (2018). Environmental DNA: For biodiversity research and monitoring. Oxford University Press.

Trunz, V., Lucchetti, M. A., Bénon, D., Dorchin, A., Desurmont, G. A., Kast, C., Rasmann, S., Glauser, G., and Praz, C. J. (2020). To bee or not to bee: The ‘raison d#x2019;être#x2019;of toxic secondary compounds in the pollen of Boraginaceae. Functional Ecology, 34(7):1345–1357.

Van Valen, L. (1965). Morphological variation and width of ecological niche. American Naturalist, 99(908):377–390.

Vaudo, A. D., Patch, H. M., Mortensen, D. A., Tooker, J. F., and Grozinger, C. M. (2016). Macronutrient ratios in pollen shape bumble bee (*Bombus impatiens*) foraging strategies and floral preferences. Proceedings of the National Academy of Sciences, 113(28):E4035–E4042.

Vaudo, A. D., Tooker, J. F., Grozinger, C. M., and Patch, H. M. (2015). Bee nutrition and floral resource restoration. Current Opinion in Insect Science, 10:133–141.

Vaudo, A. D., Tooker, J. F., Patch, H. M., Biddinger, D. J., Coccia, M., Crone, M. K., Fiely, M., Francis, J. S., Hines, H. M., Hodges, M., et al. (2020). Pollen protein: Lipid macronutrient ratios may guide broad patterns of bee species floral preferences. Insects, 11(2):132.

Waser, N. M. (1978). Interspecific pollen transfer and competition between co-occurring plant species. Oecologia, 36(2):223–236.

Waser, N. M. (1986). Flower constancy: definition, cause, and measurement. American Naturalist, 127(5):593–603.

Wells, H. and Rathore, R. R. (1994). Foraging ecology of the asian hive bee, *Apis cerana indica*, within artificial flower patches. Journal of Apicultural Research, 33(4):219–230.

Wild, J., Kaplan, Z., Danihelka, J., Petřík, P., Chytrý, M., Novotný, P., Rohn, M., Šulc, V., Brůna, J., Chobot, K., et al. (2019). Plant distribution data for the Czech Republic integrated in the Pladias database. Preslia, 91:1–24.

Zinger, L., Bonin, A., Alsos, I. G., Bálint, M., Bik, H., Boyer, F., Chariton, A. A., Creer, S., Coissac, E., Deagle, B. E., et al. (2019). DNA metabarcoding—need for robust experimental designs to draw sound ecological conclusions. Molecular Ecology, 28(8):1857–1862.

